# Characterisation of SpoT-disruption induced metabolic shift in bacteria

**DOI:** 10.64898/2025.12.06.692721

**Authors:** Zhi Y. Kho, Jinxin Zhao, Mei-Ling Han, Mohammad A. K. Azad, Christopher Barlow, Tony Velkov, Jian Li

## Abstract

SpoT, a pivotal stringent response regulatory enzyme, serves a dual function of both synthesizing and hydrolyzing the stress alarmones ppGpp and pppGpp, collectively known as (p)ppGpp. These alarmones act as global regulators, governing bacterial metabolic and physiological adaptation to diverse environmental stresses. Our previous investigation revealed that multidrug-resistant *Acinetobacter baumannii* notably upregulates SpoT in response to polymyxin treatment. Intriguingly, disrupting *spoT* gene enhances polymyxin killing. To comprehend SpoT metabolic regulatory function in mediating polymyxin tolerance, we conducted untargeted metabolomics, comparing metabolic perturbations upon polymyxin B (PMB) treatment between a *spoT*-disrupted *A. baumannii* mutant and its wild-type counterpart. Depletion of guanine-based purines GTP and GDP in the PMB-treated *spoT*-disrupted mutant suggests impaired (p)ppGpp hydrolysis, potentially leading to lethal accumulation of these molecules. PMB also induced more pronounced depletion of carbon sources (i.e., phosphoenolpyruvate, succinate, coenzyme A), energy metabolites (i.e., NADH biosynthesis pathway, ADP), amino acids, and the antioxidative system (i.e., glutathione, gamma-L-glutamyl-L-cysteine, (*R*)-*S*-lactoylglutathione) in the *spoT-*disrupted mutant. Interestingly, a distinctive time-dependent perturbation of fatty acyls was also observed following polymyxin treatment, with multiple fatty acyl conjugates significantly elevated in the *spoT*-disrupted mutant at 1 hour. However, this situation reversed at 4 hours, with more elevated fatty acyl groups in the wild-type compared to the *spoT*-disrupted mutant, indicating greater and more rapid PMB-induced membrane disruption in the *spoT*-disrupted mutant. Collectively, our findings suggest a potential role for SpoT in intricately coordinating *A. baumannii* energy expenditure and metabolite repertoire to ensure optimal functionality in stress tolerance and repair machineries (e.g., glutathione system, fatty acid regulation) following polymyxin treatment.

**IMPORTANCE:** Polymyxins, the last-resort antibiotics for multidrug-resistant Gram-negative bacteria, face increasing challenges due to growing tolerance. To tackle *Acinetobacter baumannii* polymyxin tolerance, we turned to SpoT, a (p)ppGpp synthetase/hydrolase protein. Our study focused on *spoT*-disrupted mutant, which exhibited heightened polymyxin antibacterial killing, yet the underlying mechanisms remained elusive. Through metabolomics, we unraveled key insights. Enhanced polymyxin killing in *spoT*-disrupted mutant likely resulted from lethal accumulation of stress alarmones (p)ppGpp, perturbations in carbon and energy metabolism, thiol-based antioxidant system depletion, and disrupted lipidic membrane repair. These findings illuminate the intricate metabolic signaling networks driving polymyxin tolerance in *A. baumannii*. By untangling pathogen-directed stress tolerance mechanisms, we deepen our understanding and identify potential therapeutic targets. Our discoveries offer new avenues to combat polymyxin resistance, paving the way for more effective treatments against superbugs. Through continued exploration, we can harness this knowledge to develop innovative strategies and overcome the challenges posed by polymyxin tolerance.

## INTRODUCTION

Antimicrobial resistance (AMR) poses a grave global medical challenge, rendering many antibiotics ineffective [1]. Worryingly, the discovery and development of new antibiotics have lagged behind the emergence of AMR [2, 3]. To address this pressing issue, the World Health Organization has identified multidrug-resistant (MDR) bacteria, including the Gram-negative pathogen *Acinetobacter baumannii*, as a “Priority 1: Critical” target for urgently needed antimicrobials [4, 5]. *A. baumannii*, a major culprit in nosocomial infections with a high mortality rate, particularly among immunocompromised patients, possesses a diverse array of molecular mechanisms that enable it to adapt and thrive amidst various environmental stresses [6–8]. Among these mechanisms is the stringent response, a sophisticated regulatory system that coordinates adaptive strategies in the face of different stress conditions, such as nutritional, oxidative, and osmotic stresses [9–14]. Central to this response is the cellular stress alarmones guanosine-3’,5’-bisdiphosphate (ppGpp) and guanosine 3’-diphosphate 5’-triphosphate (pppGpp) [15]. Collectively known as (p)ppGpp, these alarmones accumulate and serve as global regulators, directly binding to RNA polymerase and controlling a wide range of biological processes [15]. The stringent response allows stressed bacteria to redirect resources from growth and division towards stress tolerance-associated machinery, promoting bacterial survival until nutritional stress is alleviated [15–17].

Gram-negative bacteria, including *A. baumannii*, rely on the (p)ppGpp synthetase RelA and the multifunctional (p)ppGpp synthetase/hydrolase SpoT enzymes to finely regulate intracellular (p)ppGpp levels in response to diverse stress signals [13, 18]. For instance, amino acid starvation sensed by RelA, and various stressors, including carbon, iron, phosphate, and fatty acid starvation, sensed by SpoT [18–21]. Intricate governance of intracellular (p)ppGpp levels is vital for bacterial cell physiology, as excessive accumulation of (p)ppGpp can lead to growth inhibition and even bacterial cell death [21]. Therefore, the hydrolytic activity of SpoT plays a crucial role in (p)ppGpp-mediated signaling, particularly since it is the sole enzyme responsible for (p)ppGpp hydrolysis in organisms like [22]. However, it remains unknown whether other enzymes in *A. baumannii* are involved in (p)ppGpp hydrolysis. Despite the growing interest in (p)ppGpp-mediated virulence, antibiotic tolerance, and persistence, the precise role of SpoT in bacterial antibiotic tolerance remains largely unexplored due to the predominant use of *relA*-disrupted or (p)ppGpp-null (*relA* and *spoT* double gene-knockout) strains in existing studies [11–13, 23, 24]. In our recent study, we were the first to demonstrate the upregulation of RelA and SpoT proteins in *A. baumannii* upon polymyxin treatment, and the significant enhancement of polymyxin antibacterial killing with *spoT* disruption both *in vitro* and *in vivo* [25]. As a result, we hypothesized that a *spoT*-disrupted mutant would display distinct metabolic perturbations compared to the wild-type following polymyxin treatment, and we anticipated that the metabolomics signatures would provide pioneering insights into the mechanisms underlying the reduced polymyxin tolerance observed in *spoT*-disrupted *A. baumannii*.

In this study, we unearthed distinctive metabolic disruptions in guanine-based purines, nicotinamide adenine dinucleotide (NAD) biosynthesis, carbon utilization, amino acid metabolism, and fatty acyl metabolism in a *spoT*-disrupted mutant compared to the wild-type strain upon exposure to polymyxin B (PMB). Our findings suggest that SpoT plays a pivotal role in orchestrating *A. baumannii* energy utilization and metabolite repertoire, thereby ensuring efficient functionality of stress tolerance and repair machineries (such as the glutathione antioxidant system and lipidic membrane repair) to effectively counteract the stress induced by polymyxins.

## RESULTS

### Global metabolomic changes in *A. baumannii spoT*-disrupted mutant

Our untargeted metabolomics analysis unveiled a total of 2,298 putatively identified metabolites in each *A. baumannii* AB5075 wild-type (WT) and *spoT*-disrupted transposon mutant (*spoT*::tn) experimental group (**Table S1**). At 1 hour (h) of PMB treatment, principal component analysis (PCA) and analysis of variance (ANOVA) results demonstrated that the 2 mg/L PMB-treated *spoT*::tn mutant (*spoT* [PMB] *vs. spoT*) displayed a more pronounced metabolic perturbation compared to the PMB-treated WT (WT [PMB] *vs.* WT), resulting in 322 and 185 differentially altered metabolites (DAMs; log_2_ Fold Change [FC] >1 or <-1; FDR <0.05) in the respective comparison groups (**Figs S1A–D**). However, this pattern reversed at 4 h, revealing 142 and 309 DAMs in the respective comparison groups (**Figs S1A–D**). Further analysis using Kyoto Encyclopedia of Genes and Genomes (KEGG) pathway uncovered differential metabolic regulation of lipid (fatty acyl), amino acid, carbon, nucleotide, and peptide metabolism between the *spoT*::tn and WT at both 1 and 4 h post PMB treatment (**Figs 1, S1E–H**).

**Fig 1:**
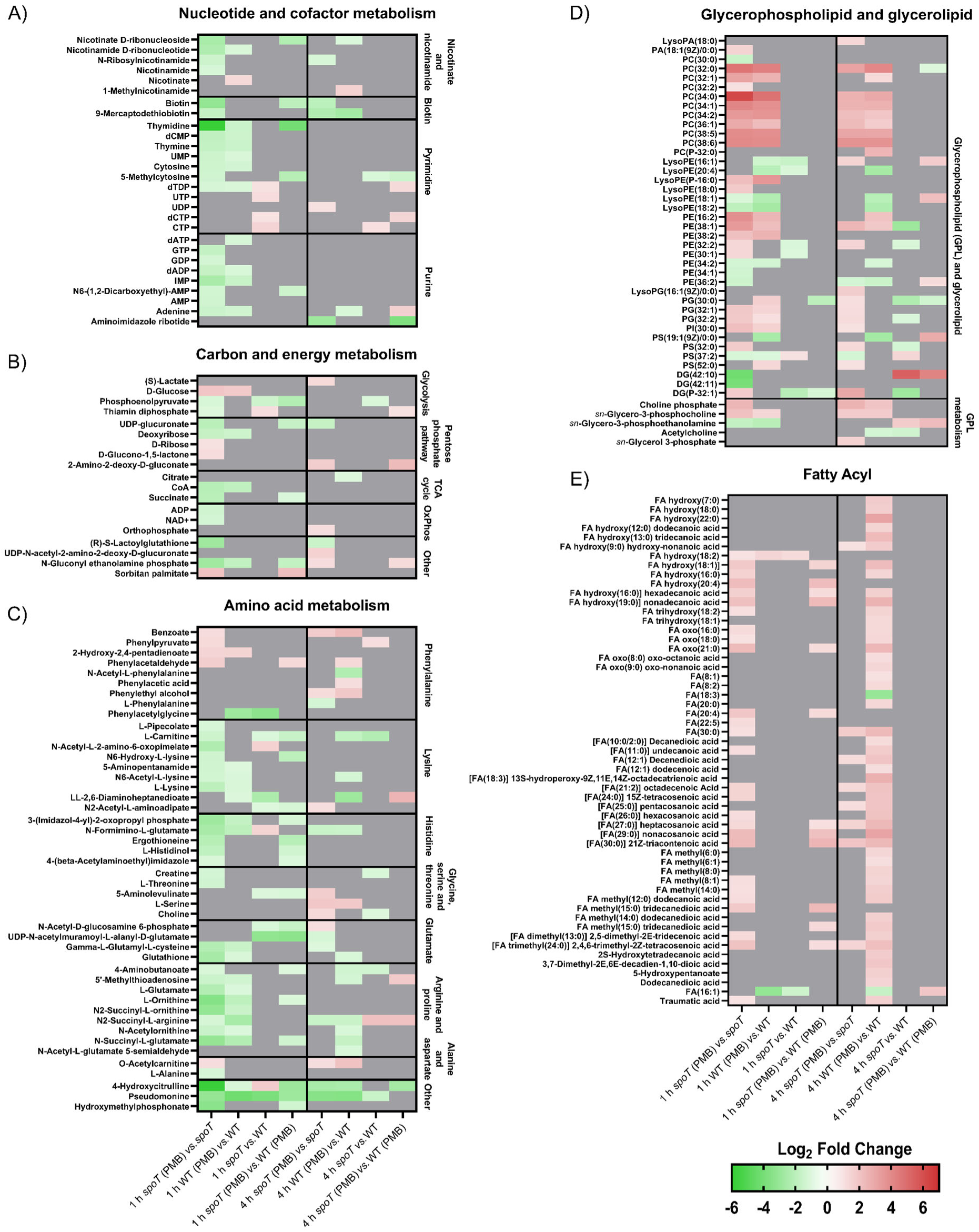
Differential metabolomic changes in the AB5075 *spoT*::tn and wild-type (WT) following 2 mg/L polymyxin B (PMB) treatment. Heatmaps delineating metabolomic changes from the categories of **A**) nucleotide and cofactor metabolism; **B**) carbon and energy metabolism; **C**) amino acid metabolism; **D**) glycerophospholipids and glycerolipids; **E**) fatty acyls in respective comparison groups across 4 h. The colour code shows log_2_ Fold Change (FC) values of differentially altered metabolites (DAMs) with log_2_FC >1 or <-1, and FDR <0.05 compared to their respective counterpart. Grey represents insignificant metabolite changes. Data are presented as the mean of four biological replicates per experimental group. OxPhos, oxidative phosphorylation; GPL, glycerophospholipid; PA, phosphatidic acid; PC, phosphatidylcholine; PE, phosphatidylethanolamine; PG, phosphatidylglycerol; PI, phosphatidylinositol; PS, phosphatidylserine; DG, diglyceride; FA, fatty acyl.

#### Polymyxin treatment reduced guanine-based purines and perturbed NAD biosynthesis in the spoT-disrupted mutant

Following 1 h of PMB treatment, the *spoT*::tn demonstrated significant reductions in guanosine triphosphate (GTP; log_2_FC, -1.73) and guanosine diphosphate (GDP; log_2_FC, -1.30) compared to its untreated control, whereas no significant changes were observed in GTP and GDP levels in the WT treatment and control groups (**Fig 2A**). Additionally, PMB treatment affected the NAD biosynthesis pathway in the *spoT*::tn mutant, resulting in reductions of intermediate metabolites such as nicotinamide, *N*-ribosylnicotinamide, nicotinamide D-ribonucleotide, nicotinate D-ribonucleoside, and NAD^+^ (log_2_FC values of -1.17, -1.52, -2.31, -2.52 and -1.28, respectively) at 1 h (**Fig 2B**). In contrast, at this time point, only nicotinate exhibited an increase (log_2_FC, 1.27) while nicotinamide D-ribonucleotide decreased (log_2_FC, -1.12) in the PMB-treated WT, with the latter showing a lesser reduction compared to the *spoT*::tn (**Fig 2B**).

**Fig 2:**
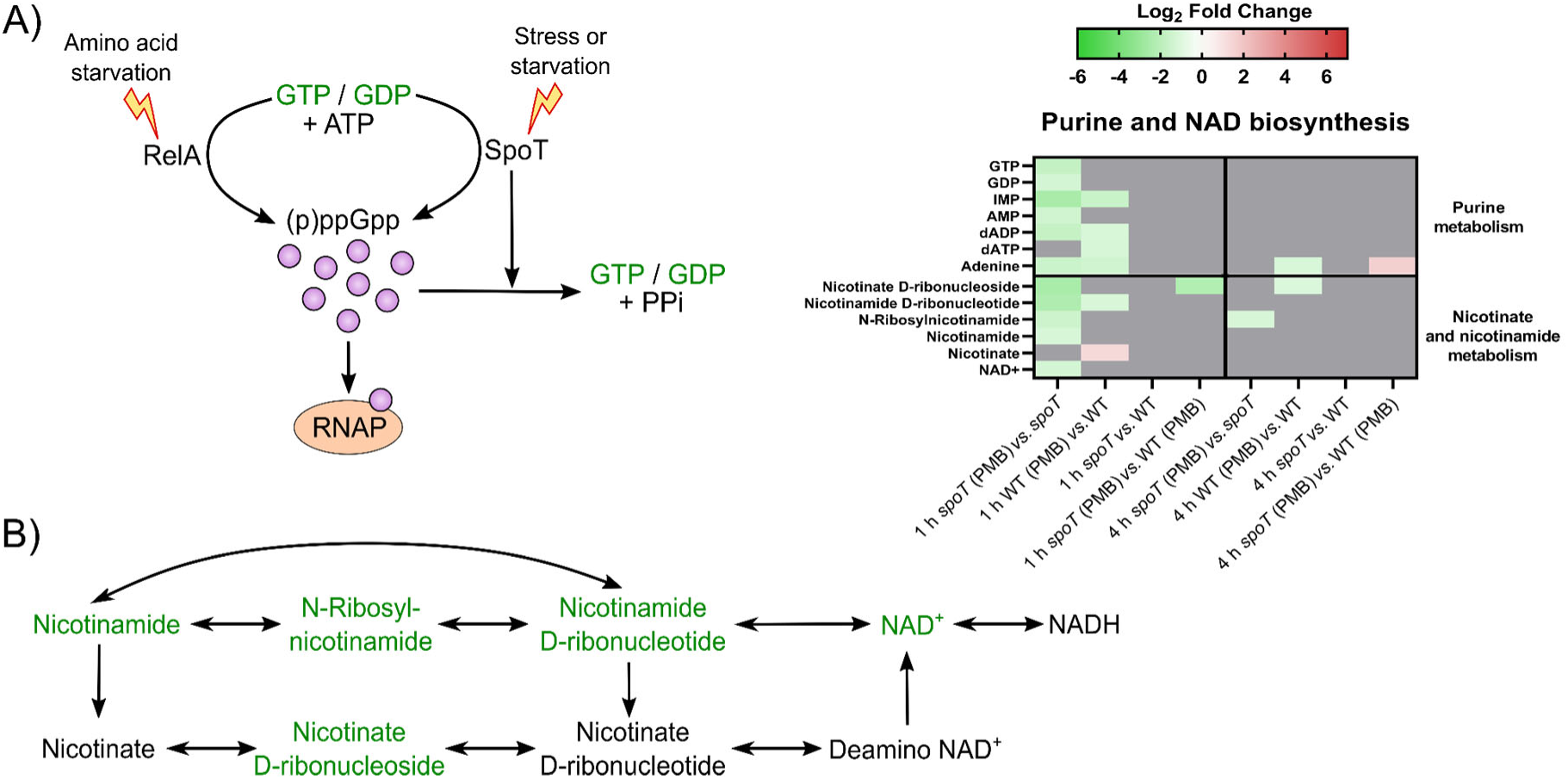
Metabolic perturbations in guanine-based purines and NAD biosynthesis in the *spoT*::tn 1 h after initiation of polymyxin B (PMB) treatment. **A**) SpoT-mediated biosynthesis and hydrolysis of (p)ppGpp. **B**) Metabolites in the NAD biosynthesis pathway of the *spoT*::tn. Differentially altered metabolites (DAMs) colored green in the schematic diagrams represent depleted metabolites in the PMB-treated *spoT*::tn at 1 h compared to its untreated control (i.e., “*spoT* [PMB] *vs. spoT*”). NAD, nicotinamide adenine dinucleotide; NADH, reduced nicotine adenine dinucleotide; RNAP, RNA polymerase.

#### Polymyxin treatment perturbed carbon and amino acid metabolism in the spoT-disrupted mutant

Metabolic substrates crucial for the tricarboxylic acid cycle and oxidative phosphorylation suffered significant perturbations in the *spoT*::tn mutant 1 h post PMB treatment. During this time, cellular glucose level was elevated (log_2_FC, 1.96), while downstream carbon and energy metabolites, namely phosphoenolpyruvate (PEP), succinate, NAD^+^, coenzyme A (CoA), and adenosine diphosphate (ADP) were all depleted (log_2_FC values of -1.18, -2.03, -1.28, -2.12 and -1.42, respectively) (**Fig 3**). Conversely, in the PMB-treated WT, glucose level was increased and CoA decreased at 1 h to similar extents as observed in the *spoT*::tn (log_2_FC values of 1.78 and -1.85, respectively), while PEP, succinate, NAD^+^ and ADP were unaffected (**Fig 3**).

**Fig 3:**
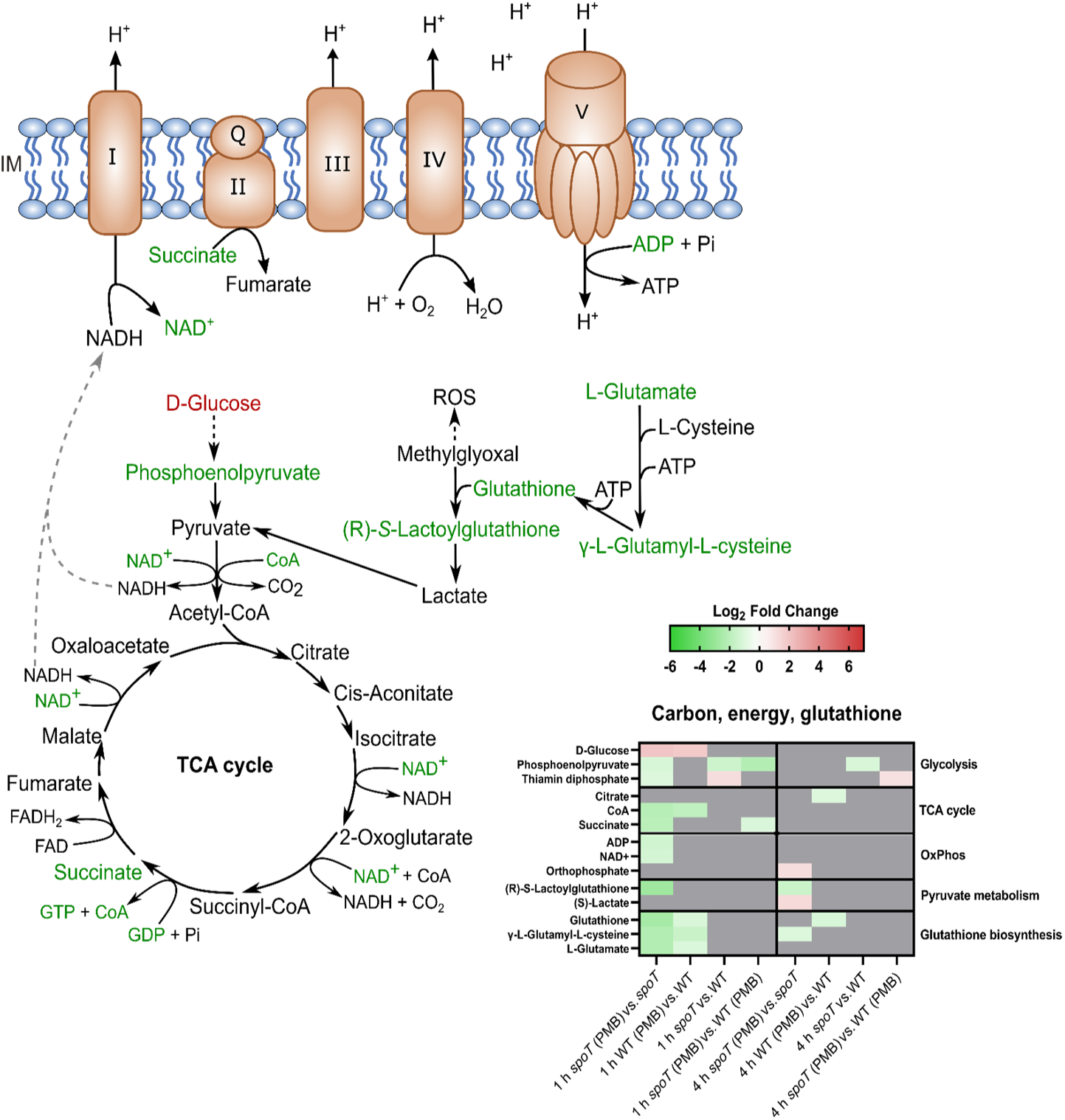
Perturbations of glutathione, carbon and energy metabolism in the *spoT*::tn 1 h after initiation of polymyxin B (PMB) treatment. In the schematic diagram, depleted (colored green) and elevated (colored red) differentially altered metabolites (DAMs) represent perturbed metabolites in the polymyxin-treated *spoT*::tn compared to its untreated control (i.e., “*spoT* [PMB] *vs. spoT*”). Data are presented as the mean of four biological replicates per experimental group. IM, inner membrane.

A strikingly greater depletion of metabolites associated with glutathione biosynthesis, namely L-glutamate, γ-L-glutamyl-L-cysteine and glutathione (GSH), was observed in the PMB-treated *spoT*::tn compared to the similarly-treated WT at 1 h (**Fig 3**). Additionally, the glutathione derivative and pyruvate metabolism intermediate, (*R*)-*S*-lactoylglutathione, a glutathione derivative and intermediate in pyruvate metabolism, showed reductions solely in the *spoT*::tn at both 1 and 4 h (log_2_FC values of -2.79 and -1.55, respectively) (**Fig 3**). Furthermore, the antioxidative histidine derivative ergothioneine (log_2_FC, -2.15) and the intermediate metabolite of histidine biosynthesis, L-histidinol (log_2_FC, -1.88), were exclusively depleted in the *spoT*::tn at 1 h (**Figs 1C, 4**). Similarly, metabolites associated with arginine biosynthesis (L-glutamate, *N*-acetylornithine and L-ornithine) and arginine metabolism (*N*^2^-succinyl-L-arginine, *N*^2^-succinyl-L-ornithine, *N*-succinyl-L-glutamate and L-glutamate) were commonly depleted by PMB treatment in both the *spoT*::tn and WT at 1 hour, but the *spoT*::tn exhibited greater depletion (except for *N*^2^-succinyl-L-arginine) (**Figs 1C, 4**). Specifically, L-ornithine and *N*^2^-succinyl-L-ornithine exhibited substantial depletion with log_2_FC values of - 3.56 and -3.16, respectively, in the *spoT*::tn, while the corresponding values in the WT were - 1.91 and -1.60. Notably, PMB treatment at 1 h led to increased levels of phenylalanine degradation pathway metabolites, particularly phenylpyruvate and phenylacetaldehyde, in the *spoT*::tn (log_2_FC values of 1.26 and 1.67, respectively) (**Fig 1C**). At 4 h, only L-phenylalanine was depleted in the *spoT*::tn (log_2_FC, -1.30), while phenylacetaldehyde (log_2_FC, 1.30) and phenylacetic acid (log_2_FC, 1.14) were elevated in the WT (**Fig 1C**).

**Fig 4:**
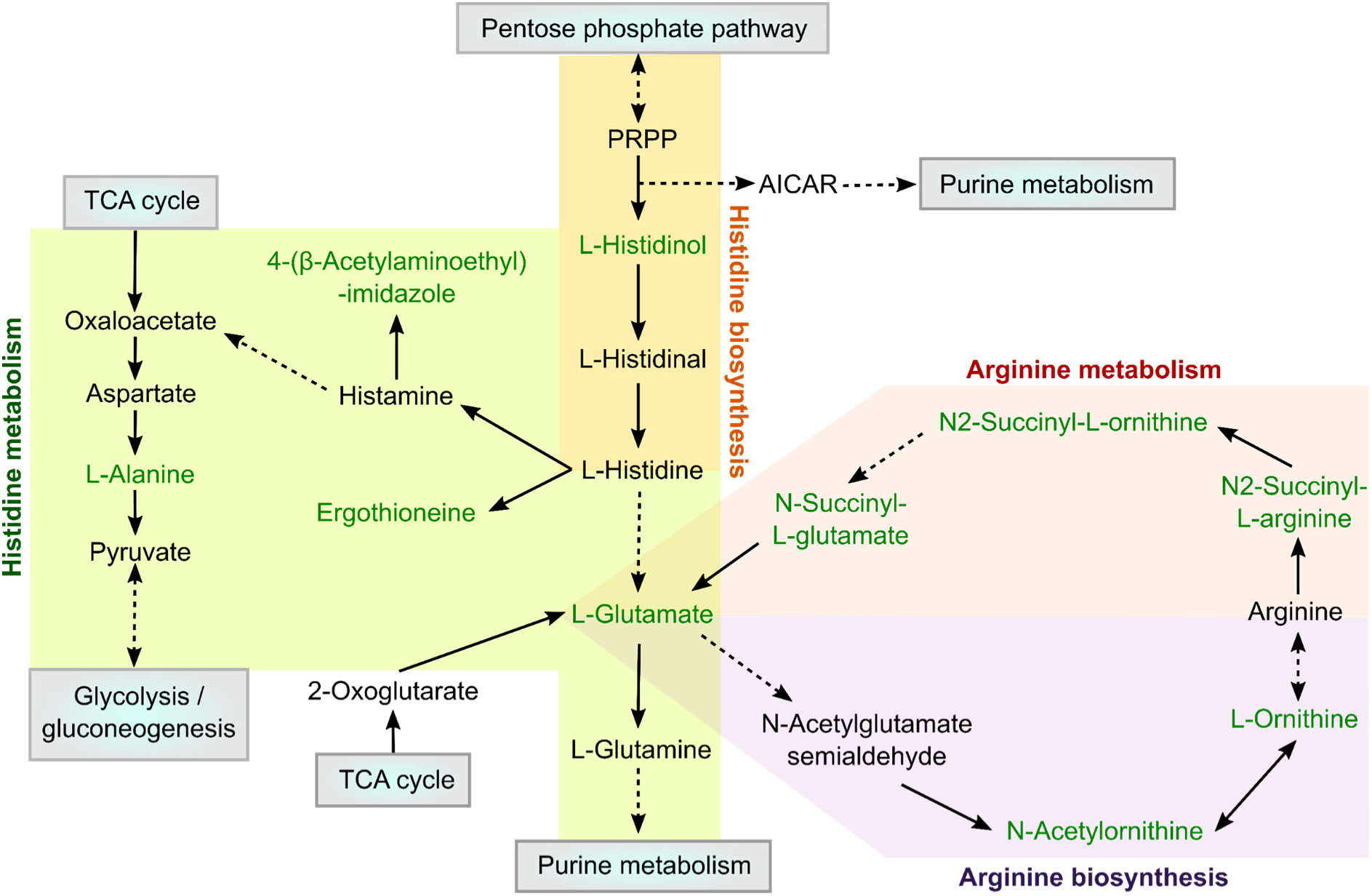
Schematic network of the perturbed amino acids in the *spoT*::tn 1 h after initiation of polymyxin treatment. Differentially altered metabolites (DAMs) colored green in schematic diagram represent significantly depleted metabolites in the polymyxin B-treated *spoT*::tn compared to its untreated control (i.e., “*spoT* [PMB] *vs. spoT*”). PRPP, 5’-phosphoribosyl-1-pyrophosphate; AICAR, 1-(5’-Phosphoribosyl)-5-amino-4-imidazolecarboxamide; TCA, tricarboxylic acid.

#### Polymyxin treatment perturbed fatty acyls and glycerophospholipids in the spoT-disrupted mutant

At 1 h post initiation of PMB treatment in the *spoT*::tn, several glycerophospholipids experienced significant elevation, including phosphatidic acid PA(18:1(9Z)/0:0), phosphatidylcholine PC(32:2), phosphatidylethanolamine PE(32:2), PE(30:1), lysoPE(18:0), and phosphatidylserine PS(32:0) (**Fig 1D**). By 4 h post PMB treatment, notable increases were observed in lysoPA(18:0), lysoPE(16:1), PE(32:2), lysoPG(16:1(9Z)/0:0), PG(30:0), PG(32:1), PG(32:2), phosphatidylinositol PI(30:0), PS(32:0) and PS(52:0) (**Fig 1D**). The PMB-treated *spoT*::tn displayed distinct perturbations in glycerolipids, with an increase in diglyceride DG(P-32:1) at 1 and 4 h (log_2_FC values of 1.74 and 2.95, respectively) and a dramatic decrease in DG(42:10) and DG(42:11) at 1 h (log_2_FC values of -4.17 and -3.95, respectively) (**Fig 1D**). Additionally, precursor metabolites for phospholipid biosynthesis showed elevated levels in the PMB-treated *spoT*::tn at both 1 h and 4 h, exemplified by choline phosphate (log_2_FC values of 2.46 and 2.59, respectively) and *sn*-glycero-3-phosphocholine (log_2_FC values of 2.11 and 1.71, respectively) (**Fig 1D**). Interestingly, *sn*-glycerol 3-phosphate (log_2_FC, 1.51), another membrane lipid precursor, exhibited exclusive elevation in the *spoT*::tn following 4 hours of PMB treatment (**Fig 1D**). Comparatively, the PMB-treated WT demonstrated similar trends, except there was no elevation in choline phosphate at 1 hour as well as *sn*-glycerol 3-phosphate at both timepoints (**Fig 1D**).

Intriguingly, a distinctive time-dependent shift in fatty acyls emerged between the *spoT*::tn and WT after PMB treatment. At 1 hour, the *spoT*::tn exhibited significantly elevated fatty acyl conjugates, unlike the mostly unchanged WT (**Fig 1E**). Surprisingly, at 4 hours, the WT revealed a greater increase in fatty acyl groups compared to the *spoT*::tn (**Fig 1E**).

#### SpoT disruption amplifies polymyxin-induced oxidative stress and membrane damage

Consistent with the metabolic alterations in redox (NAD and glutathione metabolism) and lipid biosynthesis observed in polymyxin-treated *spoT*::tn, flow cytometry revealed a marked increase in oxidative stress in the mutant. After just 15 min of polymyxin exposure, the majority of *spoT*::tn cells were CellROX Green positive (90.61 ± 2.68%), whereas the wild-type (WT) showed no significant change (**Fig 5**). Additionally, membrane depolarisation was more pronounced in polymyxin-treated *spoT*::tn, with 70.04 ± 8.60% of cells with DiBAC4(3) positive compared with untreated control, while the WT exhibited only 37.58 ± 10.01% positivity under the same treatment (**Fig 5**).

**Fig 5:**
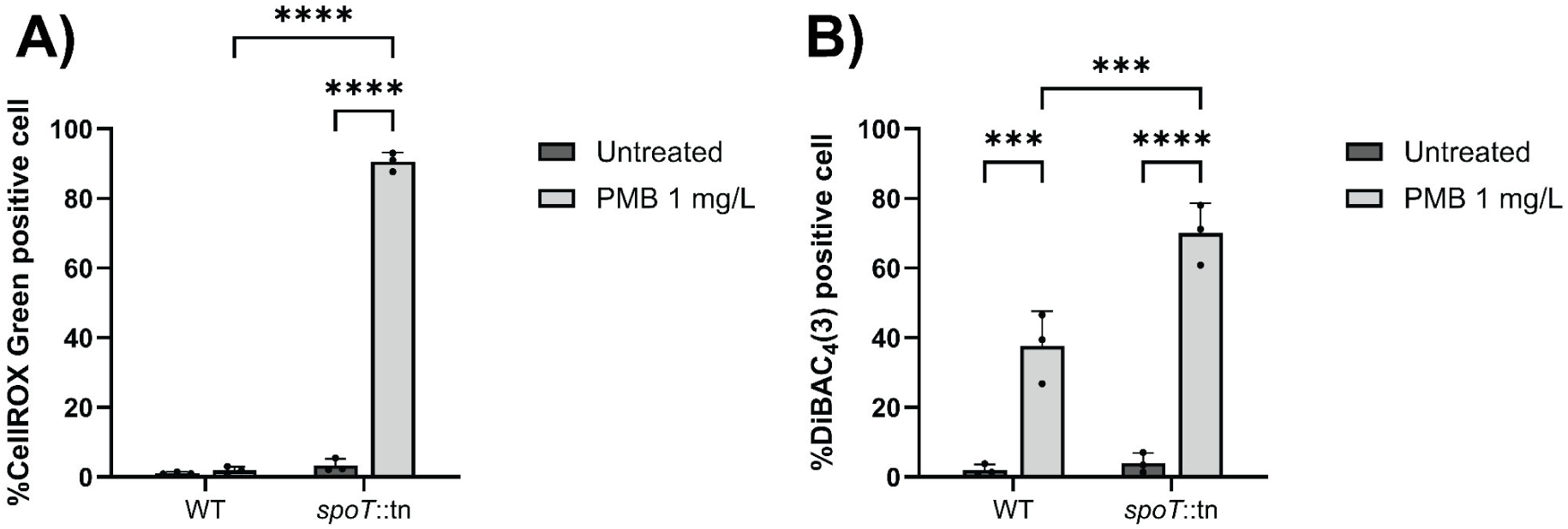
*A. baumannii spoT*::tn exhibits enhanced oxidative stress (CellROX green positive cells; **A**), and greater membrane polarity perturbation (DiBAC_4_(3) positive cells; **B**) following polymyxin B treatment at 1 mg/L for 15 min. Data in panels **A** and **B** are presented as means ± SD percentage of fluorescently stained cells from biological triplicates. Statistical significance was determined using two-way ANOVA: * p < 0.05, ** p < 0.01, *** p < 0.001, **** p < 0.0001.

## DISCUSSION

SpoT, the versatile (p)ppGpp synthetase/hydrolase, plays a pivotal role in regulating stress alarmones, making it a promising therapeutic target for multidrug-resistant bacteria. Our recent study revealed enhanced polymyxin killing in a *spoT*-disrupted *A. baumannii* mutant, showcasing the significance of (p)ppGpp-mediated stringent response [25]. Additionally, our prior proteomics investigation of *A. baumannii* wild-type reported perturbations in carbon and energy metabolism, metal homeostasis, and oxidative stress resistance machineries following polymyxin treatment, along with upregulation of (p)ppGpp regulators SpoT and RelA [25]. Based on these findings, we hypothesized that enhanced polymyxin killing of the *spoT*::tn involved aberrant signaling of (p)ppGpp, leading to the depletion of energy and precursor substrates required for *A. baumannii* stress tolerance machineries. Building on our previous proteomics findings, we utilized untargeted metabolomics to delve into the underlying mechanisms. Remarkably, we unraveled perturbations in time-dependent lipidic membrane repair and disturbed antioxidative activities in the *spoT*-disrupted mutant compared to the wild-type following PMB treatment, providing pioneering insights into role of SpoT in mediating polymyxin tolerance. This novel insight unveils potential avenues for combatting antimicrobial resistance.

SpoT senses stress signals such as starvation of carbon, fatty acid, phosphate and iron, and in response activates the stringent response by transferring a pyrophosphate from ATP to GDP or GTP to synthesize (p)ppGpp (**Fig 2A**; [19–21]). Unlike (p)ppGpp synthetase RelA, SpoT possesses a catalytic domain to hydrolyze (p)ppGpp into pyrophosphate and GDP or GTP, preventing its harmful overaccumulation that can lead to growth inhibition and bacterial cell death (**Fig 2A**; [21]). Our time-lapse live cell imaging previously showed that polymyxins induced growth inhibition and static small cells in an *A. baumannii* mutant strain harboring a transposon-disrupted *spoT* gene, suggesting (p)ppGpp overaccumulation as a likely cause [25]. This is supported by the fact that the transposon insertion site is within the hydrolytic domain of the SpoT protein of the mutant strain used (predicted using http://smart.embl-heidelberg.de/) [26]. The latest metabolomics findings in this present study unveiled significant depletion of guanine-based purines GTP and GDP in PMB-treated *spoT*::tn, indicating impaired (p)ppGpp hydrolysis and a potential toxic (p)ppGpp accumulation [21].

A remarkable finding of our study was that differential perturbations of carbon, energy and glutathione system metabolism in the *spoT*::tn compared to the wild-type shortly after commencement of PMB treatment. The increased glucose flux in both strains likely results from polymyxin-induced membrane homeostasis disturbances. However, the depletion of the downstream carbon metabolite phosphoenolpyruvate in the PMB-treated *spoT*::tn indicates an imbalance in glycolysis and failure of glucose to be converted into phosphoenolpyruvate (PEP) for entry into the tricarboxylic acid cycle, possibly due to phosphate limitation (**Fig 3**). Such an imbalance will lead to accumulation of methylglyoxal in bacterial cells [27]. Methylglyoxal is a highly reactive dicarbonyl compound which can cause oxidative stress through increased production of reactive oxygen species (ROS) and glycation with other macromolecules (e.g., lipid, protein, nucleic acid) [27, 28]. The glutathione system plays a crucial role in detoxifying such electrophiles through a two-step enzymatic reaction catalyzed by glyoxalase I and II [29, 30]. Reduced glutathione firstly conjugates with methylglyoxal to form hemithioacetal through a non-enzymatic reaction, followed by subsequent conversion into non-toxic *S*-lactoylglutathione by glyoxalase I [29, 30]. (*R*)-*S*-lactoylglutathione is then transformed into D-lactate by glyoxalase II, while glutathione is recycled [29, 30]. Prior to this step, the formed *S*-lactoylglutathione may activate the KefB and KefC potassium efflux systems, as previously demonstrated in *Escherichia coli* for protection against methylglyoxal toxicity by mediating cytoplasmic acidification [29, 31].

Our metabolomics findings reveal unique perturbations in the glutathione biosynthesis and methylglyoxal detoxification pathways in the PMB-treated *spoT*::tn (**Fig 3**). This situation could be attributed to the depletion of the essential precursor substrate L-glutamate for glutathione biosynthesis. Additionally, the observed perturbations in arginine and histidine metabolism could indirectly influence L-glutamate availability by affecting the overall metabolic network in the PMB-treated *spoT*::tn (**Fig 4**; [32–35]). Moreover, reduced flux into the tricarboxylic acid cycle and subsequent reduction in oxidative phosphorylation (**Fig 3**) could result in reduced production of energy substrates such as ATP for powering the ATP Binding Cassette (ABC) transporters (e.g., multidrug efflux, siderophore acinetobactin export, nutrient transport [36–38]), and the reducing equivalent NADH required for redox-based antioxidant systems, including glutathione system [39]. Notably, we observed a depletion of metabolites associated with the NAD+/NADH biosynthesis pathway (**Fig 2B**), indicating a substantial deprivation of NAD(P)+/NAD(P)H cellular pool in the PMB-treated *spoT*::tn. The depletion of such redox equivalents from both arms of carbon metabolism and the NAD+/NADH biosynthesis pathway could lead to enhanced perturbation of NAD(P)H-dependent redox activities in the PMB-treated *spoT*::tn, contributing to reduced tolerance to polymyxin-induced oxidative stress. This is in agreement with our previous studies involving *A. baumannii* where we demonstrated the importance of both ATP- and NADH-associated systems (e.g., acinetobactin export ABC transporter permease, thioredoxin and glutaredoxin) in stress adaptation resulting from polymyxin treatment [25, 40]. Collectively, our findings suggest that perturbations of carbon/energy metabolism and the NAD(P)H-dependent antioxidant system in *spoT*::tn likely contribute to their increased sensitivity to polymyxin-induced stress, leading to enhanced polymyxin killing.

Significant perturbations in lipid metabolism, particularly of fatty acids and phospholipids, were observed at 1 and 4 h post PMB treatment in both the *spoT*::tn and WT (**Fig 1D–E**). These findings align with the primary mode of action of the polymyxins, which involves disrupting the outer membrane, leading to increased membrane permeability and phospholipid exchange [41–44]. Intriguingly however, *spoT*::tn exhibited a distinct early increase in cellular fatty acyls at 1 h post-PMB treatment, indicating rapid membrane remodeling, which was subsequently confirmed by DiBAC_4_(3) membrane polarity staining (**Fig 5**). In contrast, the wild-type exhibited a delayed increase in fatty acyls at 4 h instead, likely reflecting slower lipid membrane perturbation and subsequent repair. Interestingly, the majority of elevated fatty acids in the *spoT*::tn were longer (i.e., 16-30 carbons) and more saturated, both of which will contribute to reduced membrane fluidity and decreased permeability to harmful substances, including polymyxins [45]. However, increased membrane rigidity will lead to growth arrest, altered cell morphogenesis and perturbed ion homeostasis [46]. Furthermore, extremely low membrane fluidity may trigger a large-scale lipid phase separation, along with segregation of membrane proteins (e.g., ATP synthase, glucose permease), resulting in disruption of essential membrane functions, as previously demonstrated in *E. coli* and *Bacillus subtilis* [46]. These differences in lipid membrane fluidity between the *spoT*::tn and wild-type likely contribute to their varying polymyxin tolerance. Additionally, fatty acids are energetically expensive molecules to synthesize, with their production requiring intricate control to match the cellular growth rate [47]. Together, the rapid increase in cellular fatty acids alongside reduced carbon metabolism in PMB-treated *spoT*::tn suggests accelerated membrane perturbation and faster energy depletion compared with the wild-type.

In conclusion, pivotal (p)ppGpp alarmones orchestrate global bacterial metabolic and physiological shifts during stress responses. Precise (p)ppGpp regulation is vital, as over-accumulation hampers growth and survival. Our study highlights SpoT, a bifunctional (p)ppGpp synthetase/hydrolase, as a potential dynamic energy regulator in response to polymyxin treatment. SpoT plays key role in ensuring an optimal metabolite repertoire, fueling stress tolerance and repair machineries (e.g., NADH-dependent antioxidants, energy-mediated transport, and lipidic membrane repair) for bacterial resilience. Together, our findings bring revolutionary revelations, supporting SpoT as promising therapeutic bullseye for boosting polymyxin killing.

## MATERIALS AND METHODS

### Bacterial isolates and antibiotics

A tetracycline-resistant AB5075-UW T26 transposon mutant with *spoT* disruption (purchased from the University of Washington [48]) and its wild-type (WT) counterpart (a wound isolate from a patient at the Walter Reed Army Medical Center [49]) were used in this study. Sterile stock of polymyxin B sulphate (PMB; Cat# 86-40302) from Betapharma (Shanghai, China) was prepared in Milli-Q™ water (Millipore, USA) and filter-sterilized using a 0.22-µm syringe filter (Sartorius, Germany). The minimum inhibitory concentrations (MICs) of polymyxin B for both the AB5075 wild-type and *spoT*::tn were determined to be 0.25 mg/L using broth microdilution as per the Clinical Laboratory Standards Institute (CLSI) guidelines (M100-Ed32). Before each experiment, frozen stocks of AB5075 wild-type and *spoT*::tn stored at -80 °C were streaked on nutrient agar and Luria-Bertani (LB) agar, respectively. LB agar was supplemented with 10 mg/L tetracycline hydrochloride for transposon mutant selection. Bacteria were then sub-cultured in cation-adjusted Mueller-Hinton broth (CaMHB) containing 20 mg Ca^2+^/L and 10 mg Mg^2+^/L and grown at 37 °C vigorous shaking (200 rpm) for 18 h.

### Metabolomics sample preparation

To prepare early logarithmic-phase (LP) broth cultures, bacteria cultured overnight in CaMHB were diluted 100-fold in fresh, pre-warmed CaMHB and grown at 37 °C for 2 h (200 rpm) to reach an optical density (OD) of ∼0.5 at 600 nm. LP cultures were then centrifuged at 3,220 × *g* and 4 °C for 20 min to pellet bacteria. The pellets were resuspended in pre-warmed Roswell Park Memorial Institute (RPMI) 1640 medium supplemented with 25 mM of 4-(2-hydroxyethyl) piperazine-1-ethanesulfonic acid (HEPES) buffer solution and 10% heat-inactivated fetal bovine serum (FBS; Lot# 15703; Bovogen, Victoria, Australia) to create a starting inoculum of ∼10^8^ CFU/mL. Four groups were prepared: untreated control for the wild-type (i) and *spoT*::tn (ii), and 2 mg/L polymyxin B (4×MIC) treated wild-type (iii) and *spoT*::tn (iv). Four biological replicates of each group were prepared from different bacterial colonies. The experimental cultures were incubated at 37 °C with shaking (200 rpm). Samples were collected at 0 (baseline), 1 and 4 h and immediately transferred to a dry ice-ethanol bath for metabolic quenching before sample normalization (OD_600_ of 0.5). Subsequently, 10 mL of each normalized sample was transferred to a 15-mL Falcon tube, and the bacterial cells were pelleted via centrifugation at 3,220 × *g* and 4 °C for 10 min prior to metabolite extraction.

The metabolite extraction method was previously described [44, 50]. Briefly, pelleted bacterial cells were washed twice with ice cold 0.9% saline to remove residual medium components and extracellular metabolites. Cell pellets were then resuspended in 500 μL of chloroform:methanol:water (1:3:1, v/v) extraction buffer containing 1 μM of four generic internal standards (CHAPS, CAPS, PIPES and TRIS; Sigma-Aldrich, Castle Hill, Australia). Samples were thrice snap-frozen in liquid nitrogen, thawed on ice, and vortexed to release intracellular metabolites. Removal of cell debris was done via centrifugation at 14,000 × *g* and 4 °C for 10 min. The particle-free supernatant (200 μL) was transferred to an injection vial for liquid chromatography-mass spectrometry (LC-MS/MS) analysis. Additionally, pooled biological quality control (PBQC) samples were prepared by mixing 10 μL of each tested sample.

### LC-MS/MS sample analysis

Metabolomics analysis was performed using a Q-Exactive Orbitrap mass spectrometer (Thermo Fisher) coupled to a Dionex high-performance liquid chromatograph (U3000 RSLC HPLC, Thermo Fisher) as described previously [44, 51]. Operation of the MS system was conducted in both positive and negative electrospray ionization mode at a resolution of 35,000 with ion detection ranging from 85 to 1,275 m/z. Metabolite samples (10 μL) underwent gradient elution through a Zwitterionic Hydrophilic Interaction Liquid Chromatography column on a polymer support (ZICpHILIC; 5 μm, polymeric, 150 × 4.6 mm; SeQuant, Merck) using LC mobile phase solvents A (20 mM ammonium carbonate) and B (acetonitrile). The elution gradient started from 80% solvent B and was reduced to 50% over 15 min with a flow rate of 0.3 mL/min, followed by a reduction to 5% solvent B over the next 3 min. Subsequently, samples were washed with 95% solvent A and 5% solvent B for 3 min, and eventually re-equilibrated with 80% mobile phase B and 20% mobile phase A for 8 min. All metabolite samples were analyzed within the same LC-MS/MS batch, and the PBQC samples were analyzed periodically throughout the batch for quality assessment. Randomized sample analysis sequence was performed to minimize operation variation. A mixture of over 250 authentic standards was included in the analysis to assist in metabolite identification.

### Data processing and bioinformatics

Metabolomics raw data was processed using mzMatch and IDEOM (http://mzmatch.sourceforge.net/ideom.php) [52]. ProteoWizard converted LC-MS/MS raw data to mzXML format [53]. Next, mzMatch was used to align and filter chromatographic peaks based on their intensity (minimum detection of 100,000 counts), relative standard deviation (RSD; <0.5 [reproducibility]), and peak shape (codadw; >0.8) [54]. Pre-processed peaks were further analyzed with IDEOM for noise filtering, putative metabolite identification, quantification, and visualization using default parameters [52]. Relative peak intensities underwent median normalization and log-transformation using MetaboAnalyst 4.0 prior to statistical analysis [55]. Global metabolomic variations among groups at different timepoints were visualized using principal component analysis (PCA) score plots (**Fig S1C – D**). One-way analysis of variance (ANOVA) determined significantly altered metabolites (DAMs) with false discovery rate (FDR) <0.05 and log_2_ fold change (FC) >1 or <-1. Pathway analysis of DAMs was conducted using KEGG (Kyoto Encyclopedia of Genes and Genomes) and BioCyc databases [32, 56]. Notably, the same metabolite could be annotated with more than one Peak ID due to repeated chromatographic peak detection in the same sample, but the Peak ID with greater relative intensity across the quadruplicates was selected as more representative of the metabolite abundance (**Table S1**).

### Assessment of oxidative stress and membrane disruption in the *spoT*::tn mutant following polymyxin exposure

Exponentially growing cultures of the AB5075 *spoT*::tn mutant and the corresponding wild-type strain were exposed to 1 mg/L polymyxin B in RPMI 1640 supplemented with 25 mM HEPES and 10% heat-inactivated FBS. Cultures were incubated at 37 °C with agitation (200 rpm). After 15 min of treatment, cells were harvested by centrifugation at 5,000 × g for 5 min at 4 °C, and the pellets were washed and resuspended in 0.9% NaCl. To assess intracellular oxidative stress and membrane depolarisation, bacterial suspensions were stained with fluorescent dye CellROX Green (Invitrogen, C10448) or DiBAC4(3) (Sigma, D8189), respectively. Flow cytometric analysis was performed using a NovoCyte flow cytometer (Agilent). Percentage of fluorescently stained cells was statistically analyzed using two-way ANOVA (GraphPad Prism 10.4.1). All experiments were conducted in biological triplicate to ensure reproducibility and statistical reliability.

## ACKNOWLEDGEMENTS

We thank Monash Proteomics and Metabolomics Platform for supporting the Mass Spectrometry-based sample processing, and Dr. Phillip Bergen for proofreading the manuscript. Z.Y.K. received financial support from Monash Graduate Scholarship.

Z.Y.K., M.-L.H. and J.L., conceptualization and experimental design. Z.Y.K., J.Z. and C.B., investigation, sample collection, mass spectrometry and data analysis. J.Z., J.L., M.-L.H., M.A.K.A., Y.Z., T.V., technical advice. Z.Y.K., manuscript writing. All authors, manuscript revision. J.L., resources. J.L., funding acquisition. J.L., M.A.K.A. and M.-L.H, supervision.

We declare no competing interests.

---

**Fig S1:**
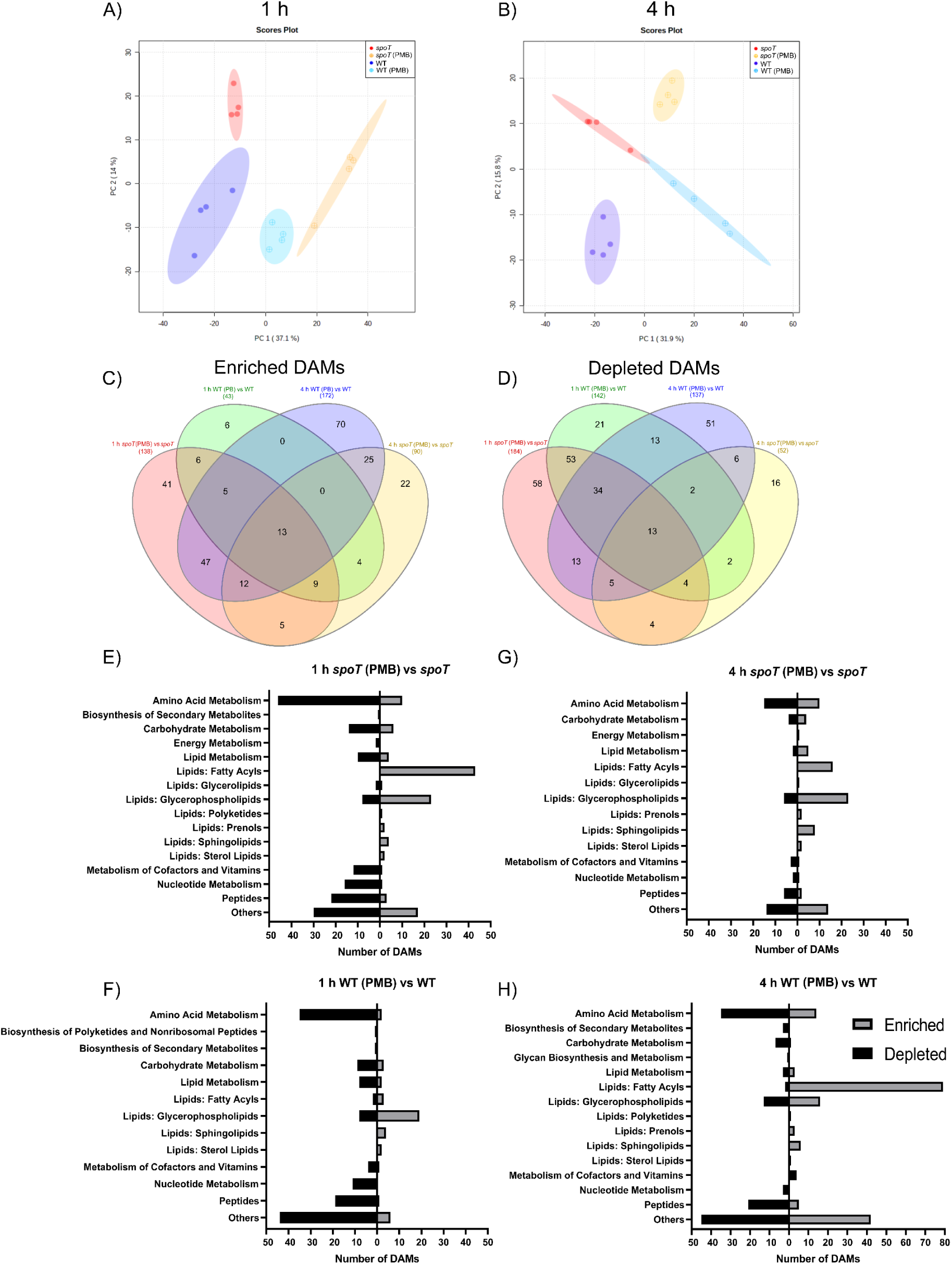
Global metabolomic changes in the *A. baumannii spoT*::tn and wild-type (WT) in the absence and presence of 2 mg/L polymyxin B (PMB) treatment. PCA score plots for 1 h (**A**) and 4 h (**B**) post PMB treatment. Venn diagram showing enriched (**C**) and depleted (**D**) differentially altered metabolites (DAMs; log_2_FC >1 or <-1, FDR <0.05) in the *spoT*::tn and WT across time. KEGG pathway analysis of DAMs of the polymyxin-treated *spoT*::tn (**E** and **F**) and WT (**G** and **H**) compared to their untreated counterpart at 1 and 4 h. Data are presented as the mean of four biological replicates per experimental group.

## List of Table

**Table S1:** List of all detected metabolites and their relative peak intensity in the absence and presence of 2 mg/L polymyxin B (PMB) treatment across time in quadruplicates. WT, wild-type *Acinetobacter baumannii* AB5075; *spoT*, *spoT*-disrupted mutant (*spoT*::tn).

